# The *Hyaloperanospora arabidopsidis* effector HaRxL77 is hypermobile between cells and manipulates host defence

**DOI:** 10.1101/2022.01.24.477405

**Authors:** Xiaokun Liu, Annalisa Bellandi, Matthew G. Johnston, Christine Faulkner

## Abstract

Multicellular organisms require dynamic communication between cells, tissues and organs to integrate responses to external and internal signals. In plants, cell-to-cell communication relies in part on plasmodesmata, which connect adjacent cells and allow the exchange of signals and resources. Upon infection by pathogens, plants act to isolate infected cells from non-infected cells by closing plasmodesmata but pathogens can suppress this defence and maintain plasmodesmata in an open state. To address the question of what a pathogen might gain from keeping plasmodesmata open, we screened effectors from the biotrophic Arabidopsis pathogen *Hyaloperonospora arabidopsidis* (*Hpa*) for the ability to move cell-to-cell via plasmodesmata in plant tissues. We quantified the mobility of cytoplasmic effectors and identified six that were hypermobile, *i*.*e*., can move further than expected for a protein of that size. Of these, HaRxL77 indirectly modifies plasmodesmatal permeability to facilitate hypermobility and suppresses the flg22-induced ROS burst, suggesting that cell-to-cell mobility of effectors allows defence manipulation ahead of the infection front. Thus, this study provides novel insights into how *Hpa* exploits plasmodesmata-mediated intercellular connectivity to promote infection, characterising a poorly explored element of plant-pathogen interaction.

**Author Summary:** During infection, pathogens secrete effectors into host cells to manipulate them for their benefit. Plant cells are connected via plasmodesmata, and pathogen effectors can use these connections to reach uninfected cells. We asked what a pathogen might gain by having effectors that can travel between cells and thus screened Arabidopsis downy mildew effectors for this capability. We found downy mildew produces many cell-to-cell mobile effectors, and that some of these can move further than we might expect. One ‘hypermobile’ effector, HaRxL77, can open plasmodesmata to allow this increased mobility, although how it does this is not clear. HaRxL77 interferes with several host defence mechanisms, suggesting that cell-to-cell mobility allows effectors to perturb host defence ahead of the infection front.

## Introduction

Plant defence against pathogens depends on the activation and deployment of plant immune responses via recognition of pathogen-derived molecules by plant immune receptors [1-3]. In the early stages of pathogen invasion, cell surface receptors recognise pathogen-associated molecular patterns (PAMPs), detecting pathogens in the apoplast and activating immune responses described as PAMP-triggered immunity (PTI) [4, 5]. Upon cellular invasion, intracellular receptors sense pathogen molecules, often proteinaceous effectors, after intracellular invasion, triggering immune responses collectively termed effector-triggered immunity (ETI) [6, 7]. ETI is highly specific and triggers strong defence responses that often establish resistance [6]. To promote disease, pathogens overcome host defence by deploying effectors that manipulate host cells to their advantage; this includes targeting and suppressing the immune signalling components of PTI and ETI [3]. Thus, the outcome of the battle between plant and pathogen can rely on which party gains control over plant immune signalling.

Most plant cells are equipped with immune receptors and signalling machinery, and thus the plant immune system can be considered as cell-autonomous. However, growing evidence supports the idea that cell-to-cell communication is also involved in plant immune responses [8, 9]. Indeed, responses such as systemic acquired resistance demonstrate that plant defence is executed in a multicellular context and intercellular communication and signalling is critical to full immune execution [10]. In plants, cell-to-cell communication can occur via plasmodesmata, the membrane-lined connections between neighbouring cells that allow movement of molecules. Molecules ranging from proteins to metabolites can traffic from cell to cell through plasmodesmata via active or passive mechanisms [11]. For example, small molecules such as metabolites can diffuse to adjacent cells via plasmodesmata [12, 13] whereas larger molecules such as transcription factors and viruses move cell to cell by employing active mechanisms that alter either the molecular structure of the cargo or the trafficking capacity of the plasmodesmata [14-16].

Our knowledge of how the plant plasmodesmata function in plant-pathogen interactions is limited, but it has become increasingly evident that plasmodesmata-mediated intercellular connectivity plays an important role in both immunity and infection. As evidence that the host regulate plasmodesmata, PAMPs including chitin and flg22 induce callose deposition and plasmodesmal closure [17, 18]. It is well established that viruses manipulate plasmodesmata to facilitate the spread of infection [19] and more recently fungal, oomycete and bacterial pathogens have been observed to exploit and target plasmodesmata. In the rice-*Magnaporthe oryzae* interaction it has been shown that the fungus delivers PWL2 and BAS1 effectors that are translocated through host plasmodesmata into neighbouring cells [20]. Additionally, effectors from *Fusarium oxysporum, Pseudomonas syringae* and *Phytophthora brassicae* have been found to directly target plasmodesmata to increase their permeability to promote infection [21-23]. These observations raise the question of what a microbe gains by maintaining symplastic connectivity in host tissues.

To address this question we have exploited downy mildew *Hyaloperonospora arabidopsidis* (*Hpa*), an obligate biotrophic oomycete pathogen of Arabidopsis [24, 25]. *Hpa* invades *Arabidopsis* cells via haustoria that are assumed to be the sites at which effectors and nutrients are exchanged between the organisms. Previous studies have determined that many oomycete effectors reside in the host cell cytoplasm [26-28], and therefore, like endogenous protiens, can possibly pass between cells through plasmodesmata to access non-infected cells. To investigate how extensive this behaviour is, and the role of cell-to-cell mobility in infection, we used a live imaging-based method to characterise the cell-to-cell mobility of a suite of *Hpa* effectors. Of these, we identified six hypermobile *Hpa* effectors. One of these, HaRxL77, confers enhanced susceptibility of the host, suppresses the flg22-triggered ROS burst and modifies plasmodesmal permeability. Characterisation of HaRxL77 provides novel insights into mechanisms by which pathogens manipulate plasmodesmata-mediated intercellular connectivity and what they might gain from this capacity.

## Results

### *Hpa* effectors expressed at 3dpi predominantly localise in the cytoplasm in planta

Previous analysis of the *Hpa* genome, led to the prediction of 475 *Hpa* genes as encoding effectors [29] and transcriptomic analysis of these predicted 475 effectors [29] revealed 87 are transcriptionally induced more than two-fold at 3dpi, an early stage of infection when we expect the cell-to-cell mobility of effectors to be relevant to establishing infection (Table S1). We assumed that mobile effector proteins would be most significantly represented amongst the cytoplasmic, soluble proteins. Thus, here our first aim was to assess these 87 effector candidates for their subcellular localization *in planta* using transient *N. benthamiana* expression. To assess the localisation of *Hpa* effectors, we cloned their sequences (with the signal peptide removed) in frame with a C-terminal GFP tag in plant expression vectors using the Golden Gate modular cloning system [30]. From 87 candidates, we successfully determined the subcellular localisation of 71 candidates in *N. benthamiana* leaves (Table S1). Of the 71 effector candidates tested, 35 (49%) localized to the cytoplasm (Fig 1 and Table S1) and we decided to restrict our mobility screen to these candidates. We further eliminated effectors that induced cell stress incompatible with live-imaging and those that had low expression levels from the screen, reducing the list of 35 cytoplasmic effectors to 19 (Table S1).

**Figure 1.**
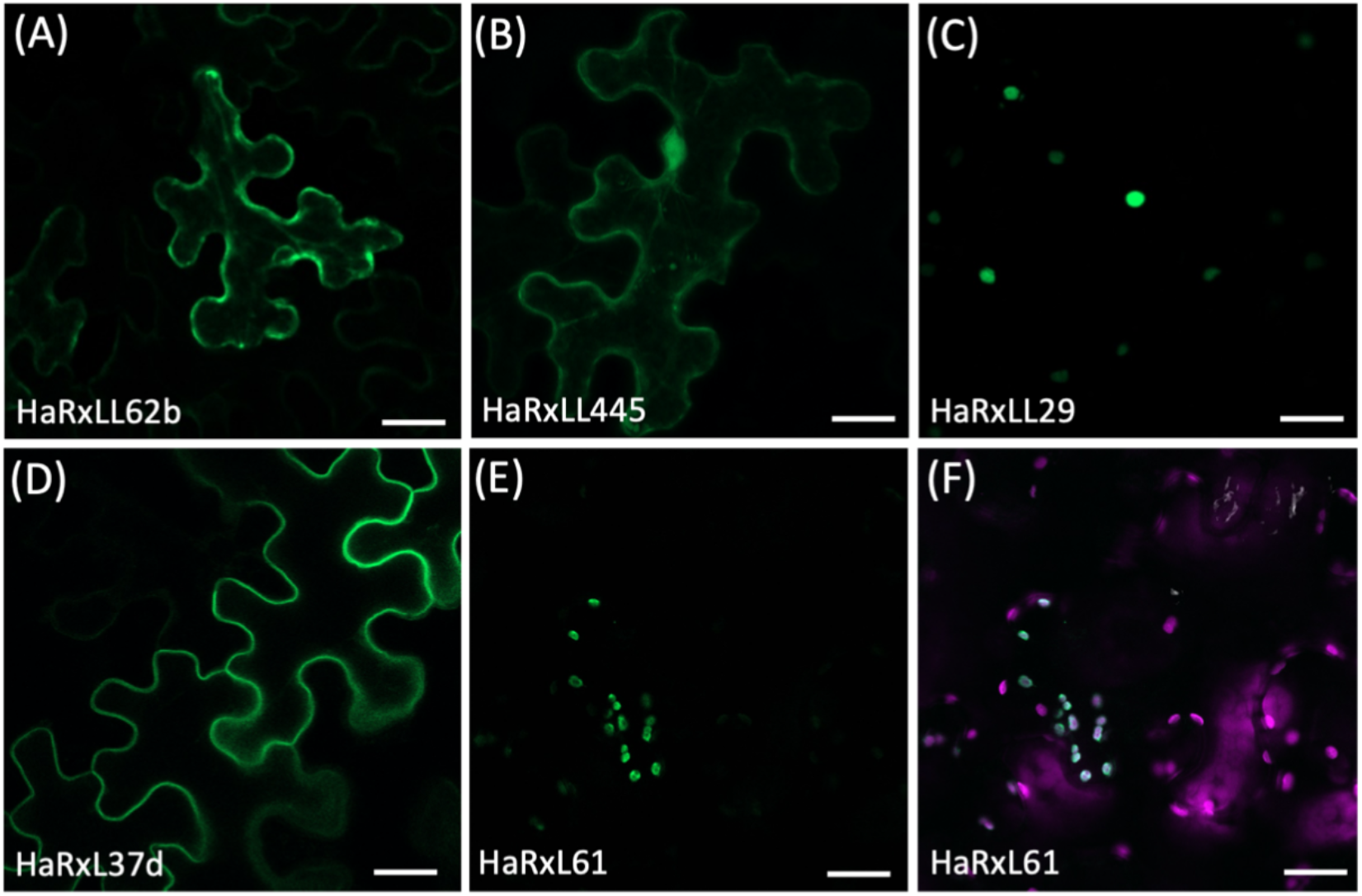
Subcellular localisations of *Hpa* effectors in *N. benthamiana* leaf cells. Effector-GFP fusion constructs were agroinfiltrated into 5-week-old *N. benthamiana* leaves, and their subcellular localisations assessed by confocal microscopy 3 days post-infiltration. Images show example micrographs of different subcellular localizations of effectors: (A) endoplasmic reticulum showing nuclear envelope; (B) cytosol; (C) nucleus; (D) plasma membrane; (E and F) chloroplast; HaRxL61-GFP fluorescence (green, E) in chloroplasts with overlay (F) of chlorophyll autofluorescence (magenta). Scale bars represent 30μm.

### Analysis of effector intercellular mobility identified a subset of hypermobile *Hpa* effectors

We previously established a live cell imaging-based method to quantitatively assay the cell-to-cell mobility of protein-GFP fusions [31]. Briefly, an effector-GFP fusion and a NLS-dTomato transformation marker were assembled into one expression vector using Golden Gate modular cloning [30] (Fig S1A). Transient expression of the two reporters in single cells of *N. benthamiana* leaves identifies the transformed cell with NLS-dTomato fluorescence and consequently infers the mobility of effector into the surrounding cells showing effector-GFP fluorescence (Fig S1B). Using this method, we screened *Hpa* effector mobility and quantified effector mobility by counting the cells showing effector-GFP fluorescence around the transformed cell [31]. Of the 19 effectors examined, 16 could move cell-to-cell and three were mmobile *in planta* (Table S1). We generated a standard curve with fluorescent proteins of known sizes to quantitively analyse effector mobility and found, of 16 mobile effectors, eight could move further than expected for a molecule of that size (were hypermobile), while one (HaRxLL468) moved less than expected (Fig 2).

**Figure 2.**
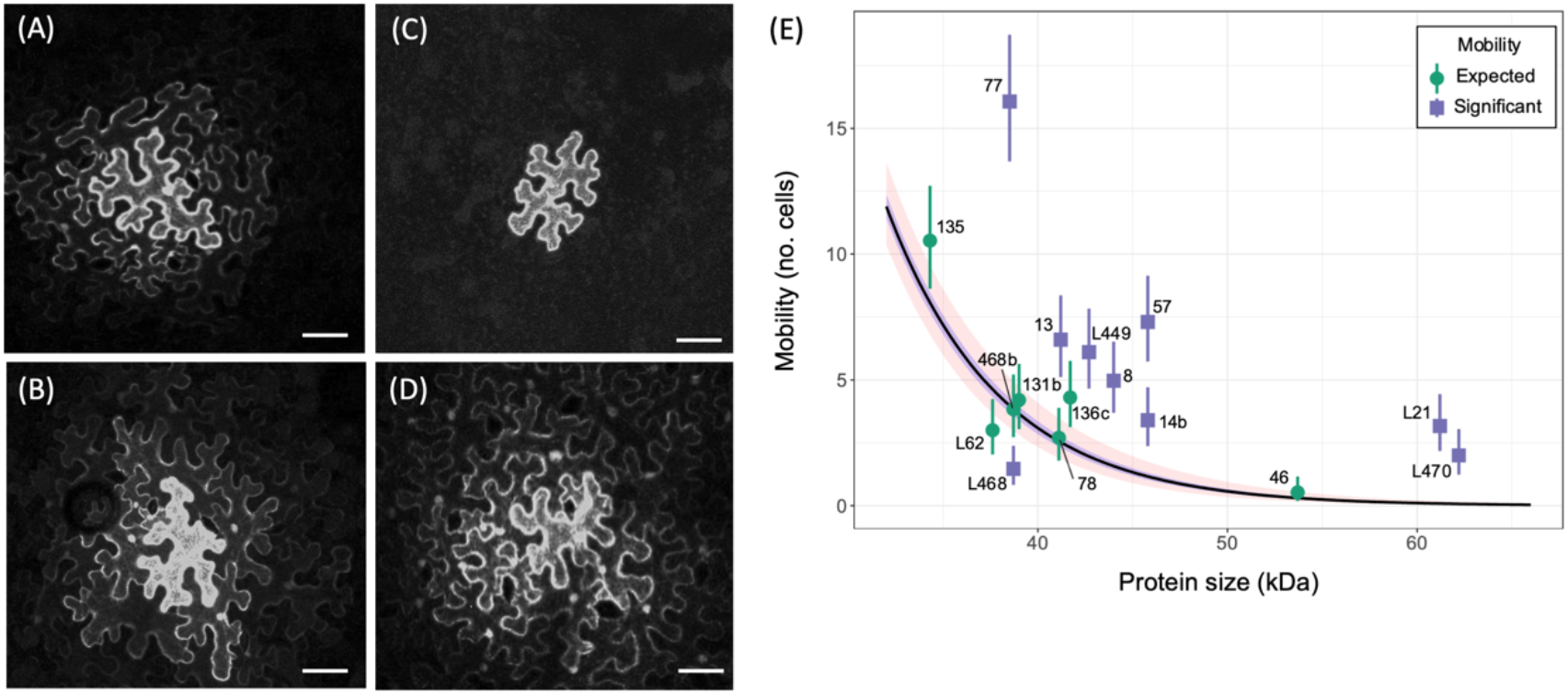
Hpa effectors are immobile, cell-to-cell mobile and cell-to-cell hypermobile. Golden Gate vectors with effector-GFP and NLS-dTomato were agroinfiltrated into 5-week-old *N. benthamiana* leaves and imaged by confocal microscopy 3 days post infiltration. Example images of mobility of effectors: (A) hypermobile (HaRxL77), (B) mobile (HaRxL135), (C) immobile (HaRxL108) and (D) cell-to-cell mobile control GFP. Scale bars represent 40 μm. (E) Size v mobility plot for 16 mobile HaRxL effectors. The curve (black line) is a size standard curve based on a quasi-poisson general linear model with a log link function and the Bonferroni corrected confidence interval of the mean (p<1×10^−5^) (red ribbon). Effectors were considered to have significant mobility (purple square) by an exact poisson test indicating the rate of movement is significantly different to the standard curve (*p* <1×10^−5^). Data points are mean ± standard error (n=30).

To exclude the possibility that the observed hypermobility of effectors arises from protein cleavage or degradation, and thus a smaller mobile protein, we extracted proteins from *N. benthamiana* tissue and detected the effector-GFP fusion by western blotting. The results showed that while some of the hypermobile effectors produced a band of small than expected size, six of eight produced a major band at the expected size. The major band from HaRxL57-GFP was smaller than expected and HaRxL14b-GFP produced two bands smaller than the expected size of the fusion protein. Thus, we concluded while HaRxL57 and HaRxL14b are likely not hypermobile, the remaining six *Hpa* effectors are most likely hypermobile (Fig S2, Table S1).

### HaRxL77 localises in both cytoplasm and plasma membrane

Of the 6 hypermobile effectors, HaRxL77 had the greatest relative mobility (Fig 2), *i*.*e*., it shows the greatest difference between the observed and the expected mobility. Further, despite some evidence of protein cleavage when expressed in *N. benthamiana*, its mobility exceeds that of GFP suggesting *bona fide* hypermobility. Thus, we chose to further characterise this effector with respect to its role in infection. Firstly, having identified that HaRxL77-GFP is mobile between cells in *N. benthamiana* leaves (Fig 2, Table S1), we aimed to confirm that HaRxL77-GFP is also mobile in Arabidopsis, the natural host of *Hpa*. Using microprojectile bombardment to mediate single cell transformation, we observed that one day after bombardment, HaRxL77-GFP moved to cells surrounding transformed cells (Fig S3). Thus, HaRxL77-GFP is a cell-to-cell mobile protein in both Arabidopsis and *N. benthamiana*.

By contrast with what we observed for HaRxL77-GFP in this study, a GFP-HaRxL77 fusion was previously reported to localise to the plasma membrane *in planta* [26],. To examine whether this difference in localisation was due to the effector being tagged on different termini, we generated two independent transgenic lines that produce HaRxL77-GFP and compared the pattern of GFP fluorescence to transgenic plants producing GFP-HaRxL77 [26]. Live imaging of leaves of these plants all showed a similar pattern of fluorescence, with both GFP-HaRxL77 and HaRxL77-GFP visible in nuclei and cytoplasmic strands as well as having a strong signal associated with the cell periphery (Fig 3A). Therefore, both N- and C-terminal fusions of GFP to HaRxL77 produce protein located in the cytoplasm.

**Figure 3.**
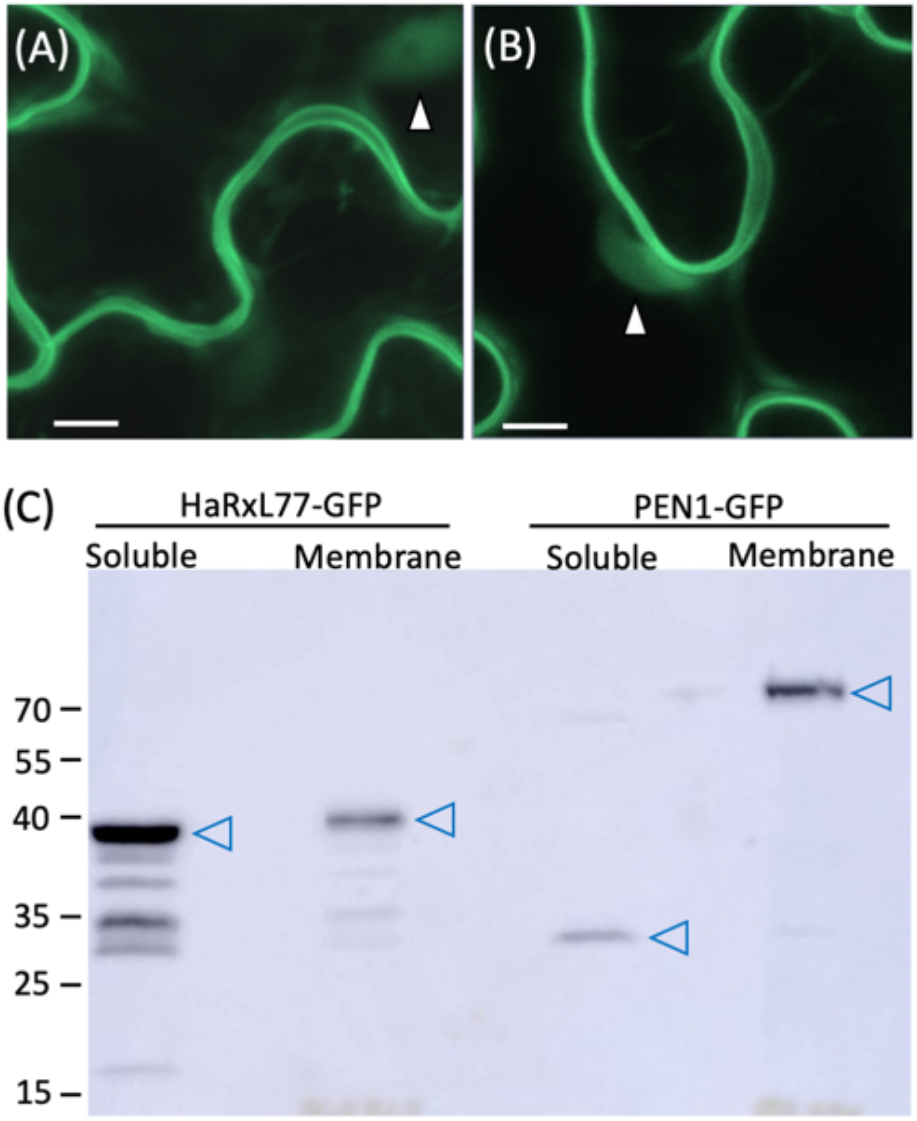
GFP fusions to HaRXL77 localise in the cytosol and the plasma membrane. Left: Confocal micrographs of cells producing (A) GFP-HaRxL77and (B) HaRxL77-GFP. White arrowheads indicate nuclei. Scale bar represents 15μm. (C) Western blot detection of HaRxL77-GFP in soluble and membrane fractions. PEN1-GFP is a plasma membrane-associated protein control. Blue arrowheads indicate the dominant bands from each fraction. Size markers are indicated on the left in kDa.

The observation of fluorescence in the nucleus and cytoplasmic strands cannot exclude the possibility that a pool of soluble, cleaved GFP populates the cytosol, while full length protein localises to the plasma membrane. To test this, we extracted proteins from the soluble and membrane fractions of *35S::HaRxL77-GFP* transgenic plants, using *35S::PEN1-GFP* transgenic plants as a control for membrane association [32]. Western blot analysis of extracts from *35S::PEN1-GFP* transgenic plants showed that a full length PEN1-GFP band is detected in the membrane fraction and a cleaved GFP band was detected in the soluble fraction (Fig 3C). By contrast and supporting the conclusion that HaRxL77-GFP has a soluble, mobile pool, full length HaRxL77-GFP bands were detected in both membrane and soluble fractions (Fig 3C).

### HaRxL77 effector confers enhanced plant susceptibility

To confirm that HaRxL77-GFP promotes virulence in the *Hpa*-Arabidopsis interaction as observed for GFP-HaRxL77 [26], we examined the effect of *in planta* production of HaRxL77-GFP on *Hpa* infection. We infected transgenic *35S::HaRxL77-GFP* plants and *35S::GFP* plants with *Hpa* and assessed *Hpa* susceptibility by counting conidiospores at 5 dpi. As observed for plants expressing *GFP-HaRxL77* [26], these data showed that plants expressing *HaRxL77-GFP* were significantly more susceptible to *Hpa* (Fig 4A), indicating HaRxL77-GFP can promote virulence of *Hpa*. Moreover, it indicates that neither N- nor C- terminal GFP fusions interfere with the virulence function of HaRxL77.

**Figure 4.**
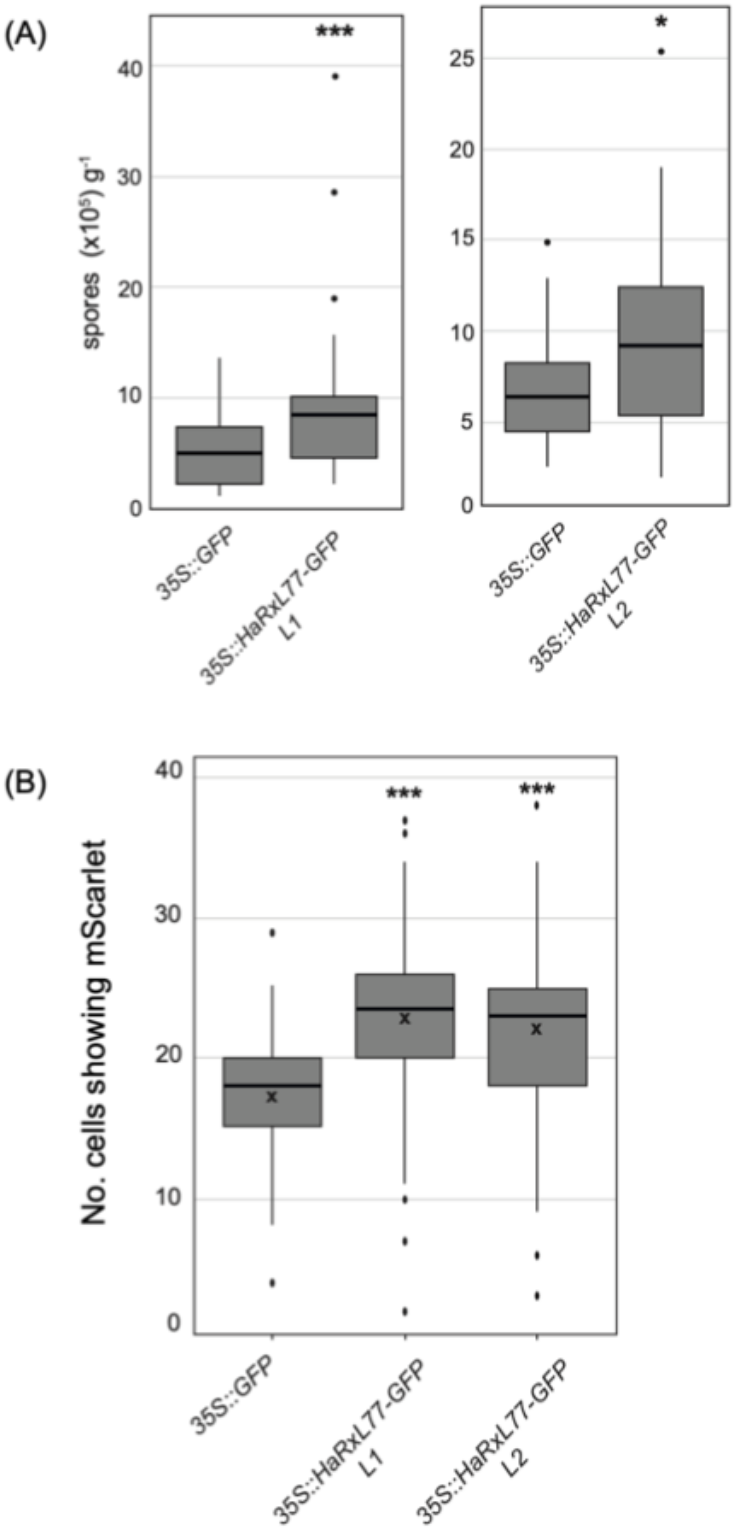
HaRxL77 confers increased virulence of Arabidopsis to *Hpa* Waco9 and increases plasmodesmal permeability. (A) Transgenic lines expressing *HaRxL77-GFP* exhibit increased susceptibility to *Hpa* Waco9. Transgenic plants were inoculated with *Hpa* Waco9 and spores were counted 5 dpi, expressed as number of conidiospores per gram fresh weight. Data was analysed by bootstrapping: \*\*\**p* < 0.001, \**p* < 0.05. Box plots: the line within the box marks the mean, the box signifies the upper and lower quartiles, and the minimum and maximum within 1.5 × interquartile range and dots indicate outliers. (B) True leaves of two transgenic *35S::HaRxL77-G*FP Arabidopsis lines and a *35S::GFP* line were bombarded with an *35S::mScarlet* encoding plasmid. The no. of cells showing mScarlet was counted for each bombardment site. Data was analysed by bootstrapping, comparing each *35S::HaRxL77-GFP* line to the *35S::GFP* control, \*\*\**p* < 0.001. Box plots: the line within the box marks the median, the cross marks the mean, the box signifies the upper and lower quartiles, and the whiskers show the minimum and maximum within 1.5 × interquartile range. Dots indicate outliers.

### HaRxL77 increases plasmodesmatal permeability

Molecules can traffic from cell to cell through plasmodesmata via active or passive mechanisms [11]. To understand the mechanism of the hypermobility of HaRxL77, we investigated whether plasmodesmatal permeability was altered in presence of HaRxL77. For this, we measured cell-to-cell mobility of mScarlet, quantified as number of cells showing red fluorescence one day post bombardment, in transgenic *HaRxL77-GFP* lines using a microprojectile bombardment assay. Mobility of mScarlet was quantified as number of cells showing red fluorescence one day post bombardment (Fig S4) and found that mobility of mScarlet was increased in both *HaRxL77-GFP* lines compared with the *GFP* transgenic control (Fig 4B, S5). These results suggest that HaRxL77 can increase plasmodesmal permeability, which likely explains its hypermobile behaviour, *i*.*e*., HaRxL77 moves further than expected for a protein of its size because it increases plasmodesmal trafficking capacity.

Plasmodesmal aperture is dynamically regulated by accumulation and removal of callose at neck regions of plasmodesmata [33] and reduced plasmodesmal callose is frequently correlated with increased plasmodesmal aperture and cell-to-cell flux [18, 34-36]. To investigate whether the HaRxL77-induced increase of plasmodesmal permeability is due to depletion of plasmodesmata-associated callose, we quantified plasmodesmal callose in effector-expressing lines using aniline blue staining (Fig 5A). We found that the mean intensity of aniline blue associated with plasmodesmal callose deposits is increased in the presence of HaRxL77-GFP (Fig 5B) suggesting that there is increased callose deposition at plasmodesmata. Thus, these results suggest that increased plasmodesmal permeability induced by HaRxL77 occurs via a mechanism independent of callose regulation.

**Figure 5.**
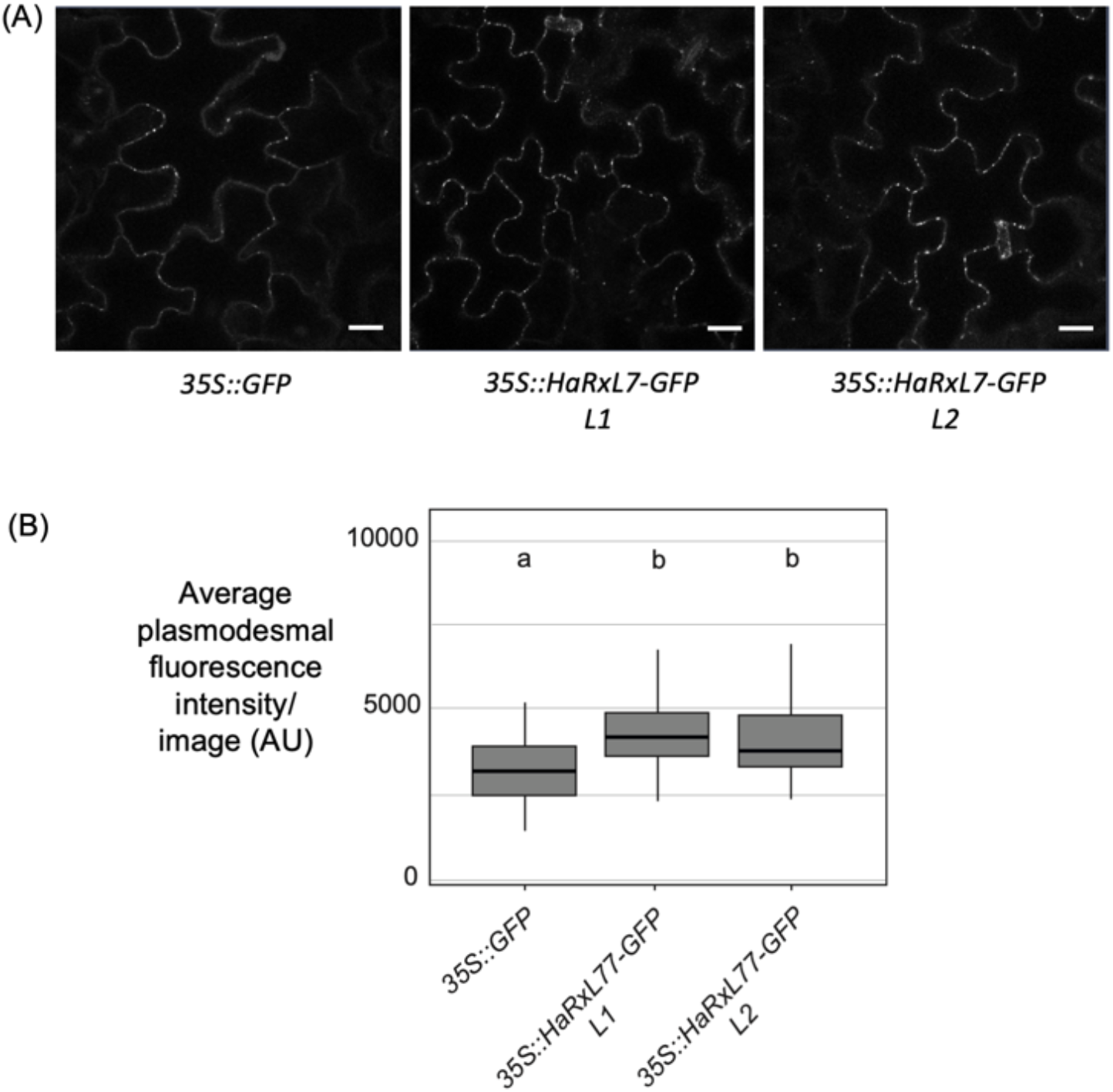
HaRxL77 lines have increased plasmodesmal callose. (A) Confocal micrographs of aniline blue staining of plasmodesmal callose in *35S::HaRxL77-GFP* transgenic lines and the control *35S::GFP* line. (B) Quantification of the average fluorescence intensity per plasmodesmal callose spot per image for *35S::HaRxL77-GFP* transgenic lines and the control *35S::GFP* line. Data was analysed by ANOVA with Tukey post-hoc analysis for multiple comparisons, different letters indicate statistically significant differences (*p* <0.005). Box plots: the line within the box marks the mean, the box signifies the upper and lower quartiles, and the whiskers show the minimum and maximum within 1.5 × interquartile range. Dots indicate outliers.

### HaRxL77 positively regulates callose deposition in the haustorial encasement

During infection, *Hpa* forms structures in host cells called haustoria that presumably act as sites of nutrient uptake [37] and effector delivery [38]. Encasement of haustoria by the host involves the deposition of materials such as callose, forming a barrier that might reduce *Hpa* access to cellular resources and block effector transit [39]. Given that we have previously observed a correlation between plasmodesmal callose and encasement callose [40], we investigated whether encasements of haustoria are altered in *HaRxL77-GFP* transgenic lines. Thus, we stained infected tissue with aniline blue and measured the width of encasements surrounding haustoria. We found that the aniline blue-stained encasement layer surrounding haustoria in *HaRxL77-GFP* transgenic lines was thicker than that in *35S::GFP* plants(Fig. 6B), while the size of haustoria was not different between lines (Fig 6A). The results further support the correlation between callose deposition in haustorial encasements and at plasmodesmata and suggests that HaRxL77 can promote both. However, by contrast with what has been seen before, the thicker encasements of haustoria in *HaRxL77-GFP* expressing lines are not correlated with greater susceptibility to *Hpa* (Fig 4A) and therefore suggests HaRxL77 might additionally targets competing immune processes.

**Figure 6.**
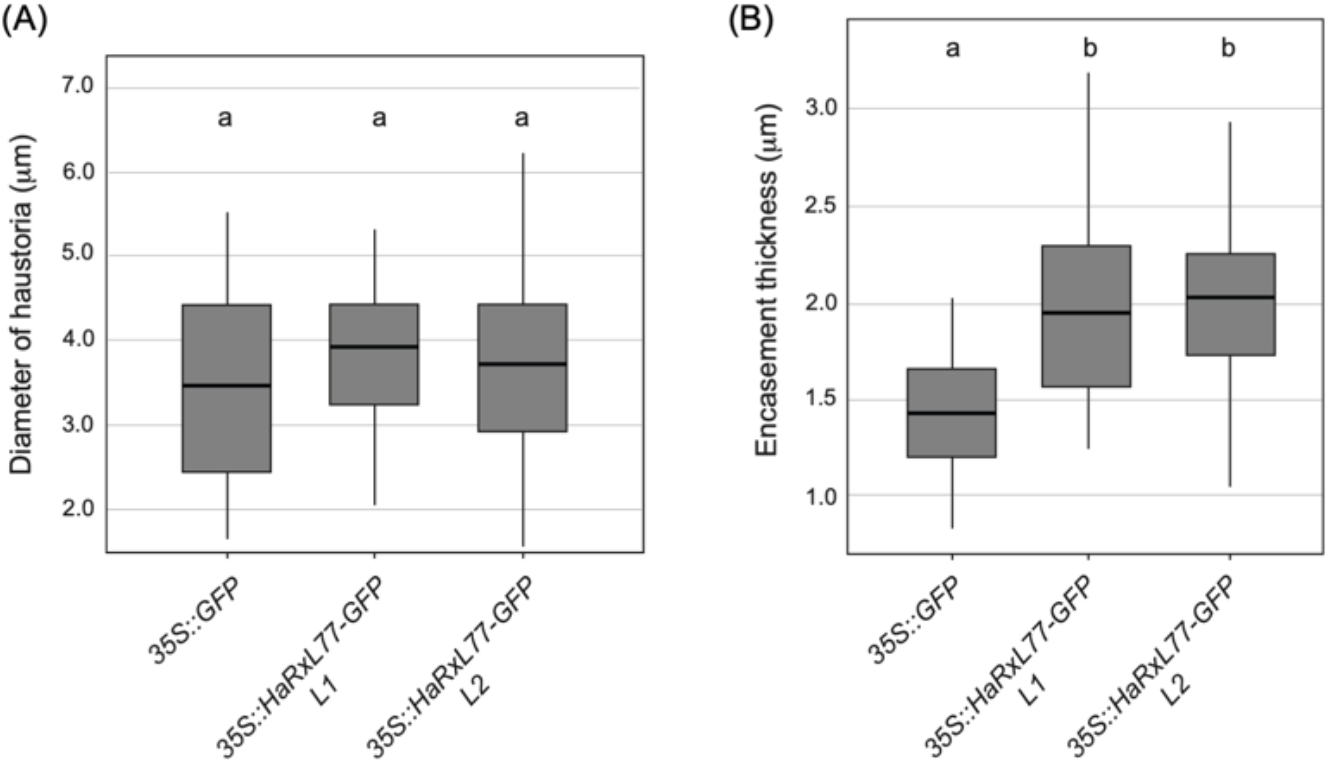
HaRxL77 positively regulates callose deposition in the haustorial encasement. 2-week old *35S::HaRxL77-GFP* seedlings were inoculated with *Hpa* Waco9 spores. At 3 dpi, aniline blue stained leaves were imaged by confocal microscopy. Haustoria (A) and encasement (B) thickness were measured in *35S::HaRxL77-GFP* transgenic lines and a *35S::GFP* control, showing that there is no difference between haustoria size in each line but that in HaRxL77-GFP transgenics the encasement is thicker. Data was analysed by ANOVA with Tukey post-hoc analysis for multiple comparisons, different letters indicate statistically significant differences (*p* <0.005). Box plots: the line within the box marks the mean, the box signifies the upper and lower quartiles, and the whiskers show the minimum and maximum within 1.5 × interquartile range. Dots indicate outliers.

Callose deposition occurs at multiple places in the cell during immune responses and so we also examined callose deposition across the cell wall during *Hpa* infection. We stained infected tissue with aniline blue and quantified the fluorescence intensity in the walls of infected and uninfected cells. In Col-0 we observed that aniline blue fluorescence is more intense in the walls of infected cells, and thus they contain more callose (Fig S6). This was also true for both *HaRxL77-GFP* transgenic lines, with the intensity of fluorescence in walls of both infected and uninfected cells comparable to that seen in Col-0 plants (Fig S6). Therefore, unlike in encasements and plasmodesmata, HaRxL77 does not enhances measurable callose deposition in walls of infected cells.

### HaRxL77 suppresses the flg22-triggered ROS burst

Rapid and transient production of reactive oxygen species (ROS) is a well-characterized response to cell surface-mediated recognition of microbes. PAMPs such as the peptides flg22 and elf18, and the carbohydrate chitin, bind receptors to activate signalling cascades that produce a burst of ROS via NADPH oxidases. To explore possible role of HaRxL77 in other immune signalling pathways, we investigated whether flg22-induced ROS burst was altered in leaves of *HaRxL77-GFP* transgenic plants. Leaf discs from 4-week-old plants were treated with flg22 and ROS production was measured by a chemiluminescence assay [41]. This showed that the flg22-induced ROS burst is significantly compromised in each *HaRxL77* line relative to *GFP* transgenic plants (Fig 7). Thus, HaRxL77 inhibits the microbe-triggered ROS burst in addition to its positive regulation of plasmodesmata and haustoria-associated callose synthesis.

**Figure 7.**
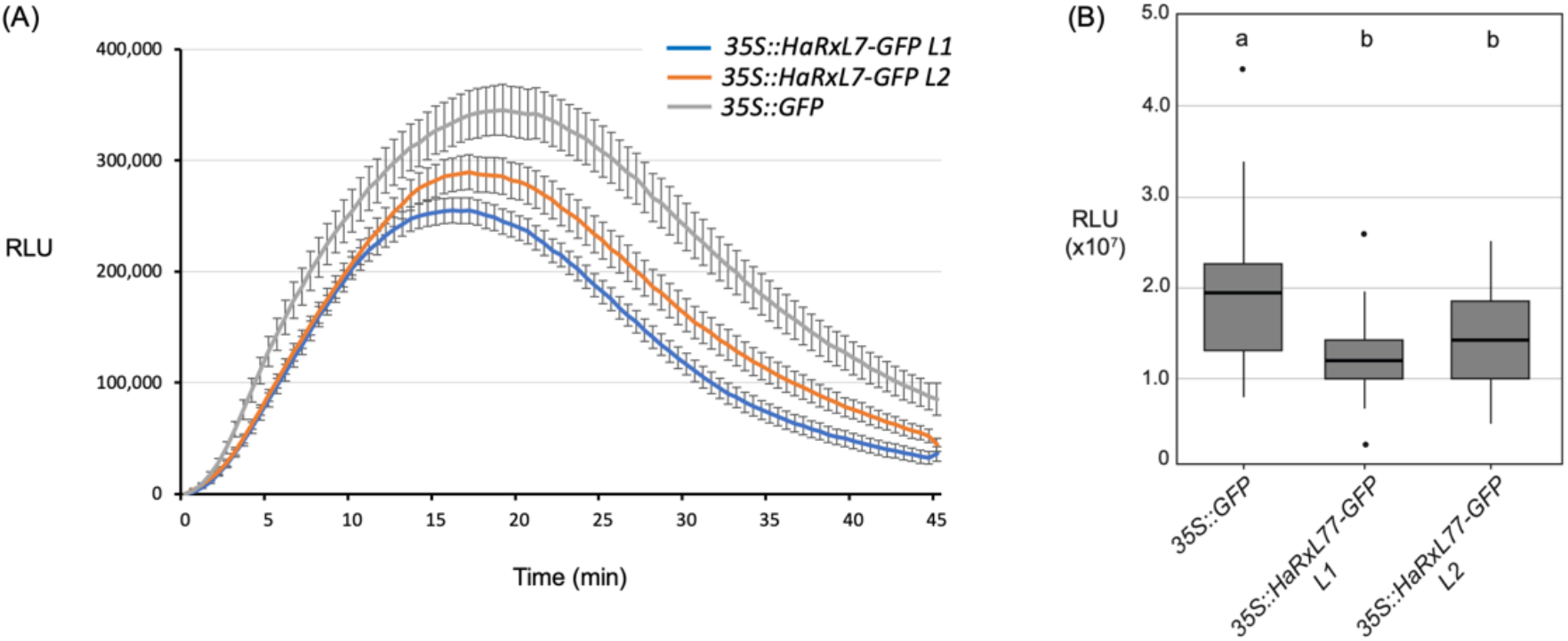
The flg22-induced reactive oxygen species (ROS) burst is compromised in the presence of HaRxL77. (A) ROS production (in Relative Light Units, RLU) was monitored in leaf discs of 5-week-old transgenic plants expressing *35S::HaRxL77-GFP* or *35S::GFP* after treatment with 100nm flg22. Error bars represent the SE of measurements from 24 leaf discs. (B) Total RLU produced within 45 mins of flg22 treatment. Data was analysed by ANOVA with Tukey post-hoc analysis for multiple comparisons, different letters indicate statistically significant differences (*p* <0.005). Box plots: the line within the box marks the mean, the box signifies the upper and lower quartiles, and the whiskers show the minimum and maximum within 1.5 × interquartile range. Dots indicate outliers.

## Discussion

A comprehensive and precise understanding of plant-pathogen interactions requires knowledge of the dynamics of molecular interactions in a spatial and temporal context. With respect to the execution of immune responses, plants have been observed to isolate cells that perceive a microbe threat by closing plasmodesmata [17]. By contrast, this cell isolation is counteracted by compatible pathogens to establish an infection presumably by action of effectors that target and suppress this response [21-23]. This observation suggests that pathogens have something to gain by maintaining plasmodesmal connectivity in the host and allowing the passage of effectors in the host tissue aids infection.

To explore this possibility, we screened effectors from the biotrophic Arabidopsis pathogen *Hpa* for intercellular mobility. From observations of endogenous mobile proteins, we expected that cell-to-cell mobile effectors would be identified from amongst cytoplasmic proteins [16, 42, 43] and therefore first identified cytoplasmic effectors for mobility screening. Like we observed for the fungal Arabidopsis pathogen *Colletotrichum higginsianum*, cell-to-cell mobility is common within the predicted effector repertoire [31], with 16 of 19 cytoplasmic effectors screened able to move between cells. Further, 6 *Hpa* effectors were hypermobile (i.e., able to move further than expected for a protein of that size) suggesting that they actively exploit plasmodesmata.

HaRxL77 was found to have greatest relative mobility of all effectors screened, and localises to the plasma membrane (Caillaud *et al*., 2012) in addition to the cytoplasm (Fig 3). We assumed that mobility of HaRxL77 is associated with the soluble pool of protein, but the *Pseudomonas* effectors HopAF1 and HopA1 are cell-to-cell mobile and associated with the plasma membrane [44] raising the possibility that this assumption is too restrictive. While there are no known plant proteins that move cell-to-cell while anchored in the plasma membrane, effectors might have novel mechanisms by which they can move cell-to-cell via the plasma membrane.

We observed that HaRxL77 hypermobilityis correlated with a capacity to generally increase the functional plasmodesmal aperture, as seen by increased mobility of mScarlet (Fig 4B) in transgenic plants that express *HaRxL77*. Given that HaRxL77 does not associate with plasmodesmata, this raises questions regarding how it might regulate plasmodesmal aperture. Further, while the data suggests plasmodesmata are more open in plants expressing *HaRxL77*, plasmodesmal callose deposition is also increased (Fig 5) in these plants. It is well established that callose deposition generally induces plasmodesmal closure, but this does not preclude novel mechanisms by which plasmodesmal aperture might be regulated. For example, mutation of *PHLOEM UNLOADING MODULATOR* (*PLM*) enhances plasmodesmata-mediated symplastic transport while not affecting plasmodesmata density or callose deposition [45]. Further investigation of the structural and functional characteristics of plasmodesmata, as well as their frequency, in *HaRxL77-GFP* transgenic plants might uncover new regulatory features of plasmodesmata.

Encasement of haustoria of *Hpa* in host cells forms a physical barrier to presumably prevent nutrient uptake and effector delivery. We previously found that a reduction in callose in haustorial encasements in the *pdlp1,2,3* mutant is correlated with increased susceptibility to *Hpa*, and conversely that overexpression of *PDLP1* leads to a thicker encasement with more callose and increased resistance [40]. The positive correlation between the callose encasement and resistance is broken in the *pmr4* mutant which has reduced callose encasement but increased resistance due to enhanced salicylic acid signalling [46]. We observed here that HaRxL77 promotes haustorial callose encasement while reducing resistance to *Hpa* suggesting that, like PMR4, HaRxL77 interacts with other elements of the immune system to enhance virulence.

The likelihood that the susceptible phenotype conferred by HaRxL77 occurs via interaction with non-callose related elements of the immune system is supported by the observation that transgenic lines that express HaRxL77 have a reduced flg22-triggered ROS burst. The ROS burst is a key immune response triggered by the activity of cell surface receptors upon perception of diverse PAMPs [5]. Membrane-localised components are involved in general ROS burst signaling machinery and often are targeted by pathogen to inhibit ROS burst [47, 48]. As there is a membrane-associated pool of HaRxL77, this effector might target plasma membrane-localised ROS signalling machinery to inhibit the ROS burst.

We have identified that *Hpa* can exploit plasmodesmata, producing effectors that can move cell-to-cell *in planta*. The observation that effectors can be both mobile and hypermobile, and modify plasmodesmal function while not directly targeting plasmodesmata, suggests that *Hpa* manipulates and exploits the host symplast in complex and indirect ways. It is well established that viruses exploit plasmodesmata to spread from cell-to-cell [19, 49], and recent studies showed that plasmodesmata are also targeted by other pathogenic microbes identifies them as a common infection target. Our observation that HaRxL77 can move cell-to-cell, and reduce the hosts defences and overall resistance, expands understanding of these observations to suggest that cell-to-cell mobile effectors enable *Hpa* to gain an advantage in infection by suppressing defence in non-infected cells.

## Methods

### Plant materials and growth conditions

*Arabidopsis thaliana* were grown on soil or on MS medium with 10 h light and 14 h dark at 22 °C unless stated otherwise. For stable transformation and selection of homozygous lines, *Arabidopsis* were grown on soil under the long day conditions with 16 h light and 8 h dark at 22 °C. *Nicotiana benthamiana* plants were grown under long day conditions at 23 °C with 16 h light and 8 h dark.

### DNA constructs

*Hpa* predicted effector sequences were obtained from the *Hpa* genome [29]. Signal peptide of the effectors were predicted by SigalP2.0 and were removed from coding sequences. Coding sequences were domesticated to remove BpiI and BsaI sites and synthesised as Golden Gate-compatible Level 0 modules. Modules were assembled as plant expression constructs using the Golden Gate cloning method [30] and all module information is in Table S1. Effector candidates were tagged with C-terminal GFP and expressed from the CaMV 35S promoter. For live cell imaging-based mobility quantification, the dual cassette constructs were assembled as shown in Fig S1 [31].

### Transient and stable expression of genes in *planta*

For transient expression in *N. benthamiana, Agrobacterium tumefaciens* (GV3101) carrying binary plasmids were infiltrated into *N. benthamiana* leaves with OD_600nm_ = 0.5 to assess subcellular localisation and with OD_600nm_ = 1.0 × 10^−5^ to quantify intercellular mobility. Transgenic Arabidopsis plants of the Col-0 ecotype were transformed by floral dipping [50]. Transgenic offspring were selected on MS medium containing 20µM L-Phosphinothricin and confirmed by confocal microscopy, genotyping PCR and western blotting. Plants homozygous for the transgene were used for experiments.

### Pathogen assays

*Hpa* growth and infection assays were performed with the Waco9 isolate, and modified from Caillaud *et al*.[40]. For infection, 10 day old *Arabidopsis* seedlings were spray-inoculated to saturation with a spore suspension of 5×10^4^ spores/ml. Inoculated plants were kept in a growth cabinet at 16°C for 3 to 6 days with 10 h light and 14 h dark. To evaluate conidiospore production, 10 pools of 3 seedlings were harvested in 1 mL of water for each line. After vortexing, the number of liberated spores was determined with a haemocytometer as described [51]. Data was collected from three independent experiments and analysed using ANOVA.

### Confocal Microscopy

For determination of the subcellular localization of effectors in *N. benthamiana* and in Arabidopsis transgenic lines, leaf samples were mounted in water and analysed on a confocal microscope (Zeiss LSM800) using the following excitation/emission wavelengths: GFP, 488/509-530 nm; RFP, 561/600-640nm. For the mobility assay, leaf tissue was imaged with a 20x water-dipping objective (W N-ACHROPLAN 20x/0.5; Zeiss).

### Quantification of cell-to cell mobility images and data analysis

Mobility quantification was performed as in [31]. Briefly, the number of fluorescent cells around the transformed cell, identified by NLS-dTomato fluorescence, was counted for each effector-GFP fusion. The data obtained was analysed in R (R Core Team, 2020). Firstly, mobility data from four markers (GFP: 26 kDa; YFPC-GFP: 37 kDa; YFPN-GFP: 45 kDa; and 2xGFP: 52 kDa) was used to generate the standard curve using a quasi-Poisson general linear model with a log link function and a Bonferroni corrected p-value < 1x 10^−5^. The mobility of each effector candidate was tested against the null hypothesis that its mobility is as predicted by the standard curve with the equivalent of a t-test for Poisson distributions (*p* < 1x 10^−5^).

### Protein extraction and Western blotting

Protein fractionation was modified from Chen. *et al*. [52]. Leaves samples were homogenized in liquid nitrogen and incubated with cytosol extraction buffer (50 mM Tris-Cl pH 7.5, 150 mM NaCl, 2 mM DTT, protease inhibitor cocktail (Sigma), 0% NP-40 (Sigma), 0% glycerol) and spun at 4 °C at 20000 g for 30 min. The supernatant was collected as soluble fraction. The pellets were washed with cytosol extraction buffer for three times and resuspended in membrane extraction buffer (50 mM Tris-Cl pH 7.5, 150 mM NaCl, 2 mM DTT, protease inhibitor cocktail (Sigma), 0.5% NP40, 10% glycerol) and centrifuged at 4 °C at 20000 rpm for 20 mins. The supernatant was collected as membrane fraction. Portions of both fractions were then mixed with 4 × sample SDS buffer (250 mm Tris-HCl, pH 6.8, 40% glycerol, 0.02% bromophenol blue and 10% β-mercaptoethanol) and boiled at 70 °C for 10 min. The proteins were separated on 12% SDS-polyacrylamide gel and transferred to PVDF membrane (Bio-Rad). Proteins were detected with an anti-GFP antibody HRP conjugate.

### Microprojectile bombardment

Microprojectile bombardment assays were performed essentially as described [17]. 10-day-old seedlings true leaves of *Arabidopsis* lines were bombarded with 1 nm gold particles (Bio-Rad) coated with plasmid DNA, using a Biolistic PDS-1000/He particle delivery system (Bio-Rad). Bombardment sites were imaged 24 h post bombardment by confocal microscopy (Zeiss LSM800) with a 10x (EC Plan-NEOFLUAR 10× 0.3; Zeiss) or 20x dry objective (Plan-APOCHROMAT 20x/0.8; Zeiss). Data were collected from at least 2 independent bombardment events, each of which consisted of leaves from at least 3 individual plants. The median mobility (number of cells) for different lines was compared in R statistical computing language v4.0.3 by a bootstrap method [53].

### Leaf disc ROS burst assay

For measurements of flg22-triggered ROS production, leaf discs were punched from 4-week-old Arabidopsis leaves with a cork borer. Leaves discs were allowed to recover overnight in water and assayed in 20 µg/mL HRP and 6 µM L-012 in a 96-well luminometer plate (Thermo Fisher Scientific), with or without 100 nm flg22. Chemiluminescence was recorded using a Varioskan Flash (Thermo Fisher), and the luminescence emitted in the first 60 mins after elicitation was integrated, corrected for background luminescence, and used for subsequent analysis.

### Plasmodesmal callose staining, imaging and quantification

For callose staining, the true leaves were detached from 2-week-old Arabidopsis seedlings and placed on a microscope slide on a droplet of 0.5% aniline blue solution in PBS buffer (pH7.4). A coverslip was gently pressed on the leaf to allow dye penetration into the apoplastic space of leaves. The leaves were then rinsed in sterile water before mounting on a microscope slide. Images were acquired on a confocal microscope (LSM Zeiss 800) using a water immersion 63x objective. Aniline blue was exited at 405 nm and emission was collected in the 410-470 nm. For plasmodesmal callose quantification, Z-stacks were acquired in two sites of each leaf for 20 leaves for each genotype.

For automated plasmodesmal callose quantification, aniline blue signal at plasmodesmata was automatically quantified in Fiji [54] using an in-house script (https://github.com/faulknerfalcons/PD_detection). The data obtained were analysed in R [55] to obtain for the mean number, size and integrated density of the detected callose particles in each z-stack. In other words, this analysis determined the mean intensity of aniline blue staining signal per plasmodesma in each z-stack, which correlates with the levels of callose.

### Callose staining and quantification of haustorial encasement and cell walls

For the measurement of callose haustorial encasements, aniline blue staining was modified from Caillaud *et al*.[26]. True leaves were detached at 3 dpi from infected Arabidopsis seedlings and were immersed in 70% Et-OH for fixation overnight. The leaves were stained in 0.1% aniline blue solution by vacuum infiltration for 2 mins and dried on tissue paper. For cell wall callose staining, true leaves were detached at 3 days after *Hpa* infection and processed as for plasmodesmal callose staining. The leaves were mounted on microscope slides and images by confocal microscopy.

For quantification of encasement thickness, the thickness of the aniline blue stained haustoria encasement was manually measured in Fiji. For quantification of cell wall, aniline blue signal at cell wall was manually measured in Fiji. The data obtained were analysed for the mean intensity of aniline blue staining signal at cell wall.

## Funding

XL was supported by Marie Curie Individual Fellowship from European Commission (749745, “HOPESEE”). Work in the Faulkner lab is supported by the European Research Council grant (725459, “INTERCELLAR”), the Biotechnology and Biological Research Council Institute Strategic Programme ‘Plant Health’ BBS/E/J/000PR9796 and by CEPAMS-Newton Fund. AB was supported by the Norwich Research Park Doctoral Training Programme and MGJ was supported by the John Innes Foundation Rotation PhD programme.

## Acknowledgements

We would like to thank Prof. Jonathan Jones from The Sainsbury Laboratory for kindly providing transgenic GFP-HaRxL77 lines seeds; Dr. Andrew Breakspear for helping setup ROS burst measurement; Dr. Jeroen De-keijzer for discussion. The authors declare they have no conflict of interest.

## Author Contributions

Conceptualization – all authors; Data curation – XL; Formal analysis – XL, AB, MGJ; Funding acquisition – XL, CF; Investigation – XL; Methodology – XL, AB, MGJ; Project administration – XL, CF; Resources – XL; Software – AB, MGJ; Supervision – CF; Visualization – XL, AB, CF; Writing – original draft – XL, CF; Writing – review & editing – XL, AB, MGJ, CF.

## Competing Interests

The authors have no competing interests

## Notes

### Competing Interest Statement

The authors have declared no competing interest.

