## Supplemental Figures for "The *Hyaloperanospora arabidopsidis* effector HaRxL77 is hypermobile between cells and manipulates host defence"

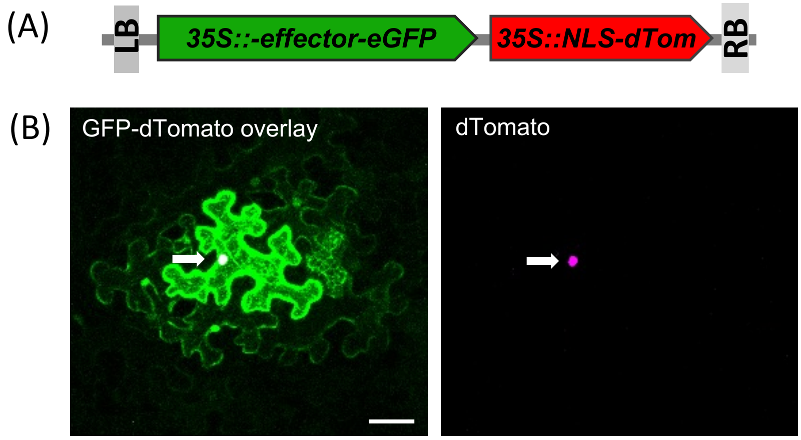


**Figure S1. Dual expression of effector-GFP and a cell transformation marker in single cells allows for detection of cell-to-cell mobility.** (A) Schematic cartoon of dual expression vector containing a 35S::effector-eGFP expression module and a 35S::NLS-dTomato expression module. (B) Live imaging of an example vector expressed in *N. benthamiana* leaves. The transformed cell is identified by magenta fluorescence in the cell nucleus (right panel, arrow). Effector-GFP fluorescence (left panel) can be detected in the transformed cell (arrow) as well as surrounding cells indicating cell-to-cell mobility. Scale bar represents 50µm.

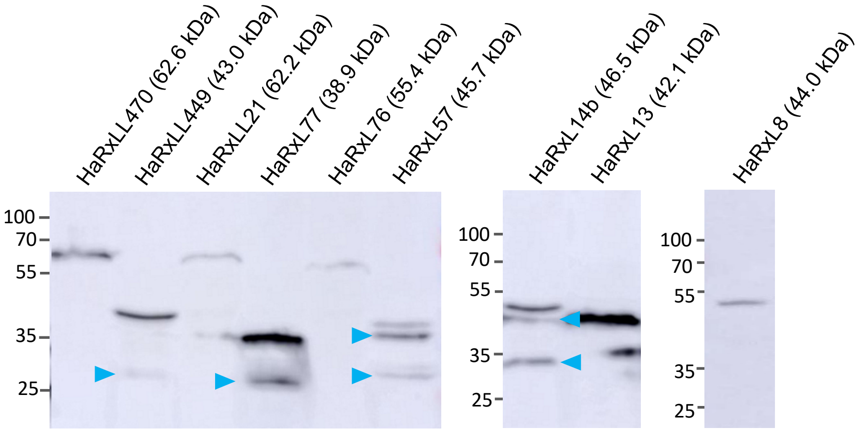


**Figure S2. Western blot analysis of hypermobile effector proteins.** Effector-GFP constructs were transiently expressed in *N. benthamiana* leaves. Total extracts were purified from leaves harvested 3 dpi and were separated on an SDS-PAGE gel. Effector-GFP fusions were detected with anti-GFP antibody (expected GFP protein fusion sizes indicated in parentheses). Blue arrowheads point to bands that suggest protein cleavage.

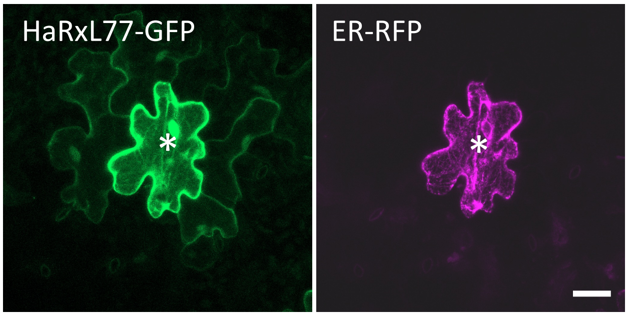


**Figure S3**. **HaRxL77-GFP is mobile in Arabidopsis leaves.** True leaves of Arabidopsis Col-0 were bombarded with gold particles coated with plasmids encoding *35S::HaRxL77-GFP* and *35S::ER-RFP*. The images were collected by confocal microscopy at 24 hours after bombardment. * indicates the transformed cell (marked by ER-RFP), and HaRxLL77-GFP fluorescence can be seen in the surrounding cells. The scale bar represents 15 μm.

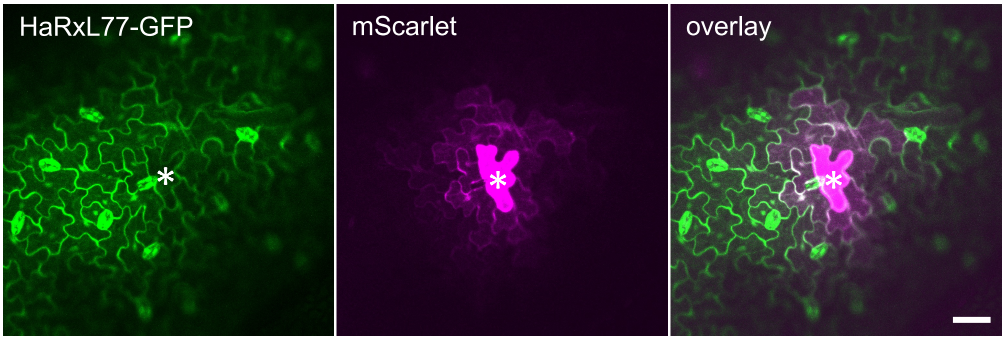


**Figure S4. Bombardment of mScarlet into leaves of *35S::HaRxL77-GFP* transgenic lines.** True leaves of 14-day-old Arabidopsis seedlings were bombarded with gold particles coated with a *35S::mScarlet* encoding plasmid. Bombardment sites were assessed by confocal microscopy at 24 h after bombardment. The transformed cell is indicated by *, the scale bar represents 50 µm .

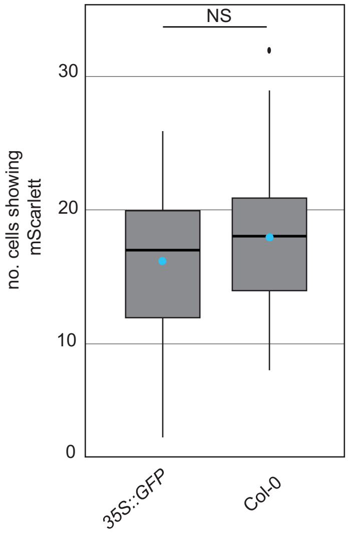


**Figure S5. GFP does not affect plasmodesmal permability.** True leaves of Arabidopsis Col-0 and a *35S::GFP* transgenic line were bombarded with gold partiles coated with a plasmid encoding *35S::mScarlet*. The no. of cells showing mScarlet was counted for each bombardment site. Data was analysed by a bootstrap method and found to be not significant (NS). Box plots: the blue dot within the box marks the mean, the box signifies the upper and lower quartiles, and the minimum and maximum within 1.5 × interquartile range.

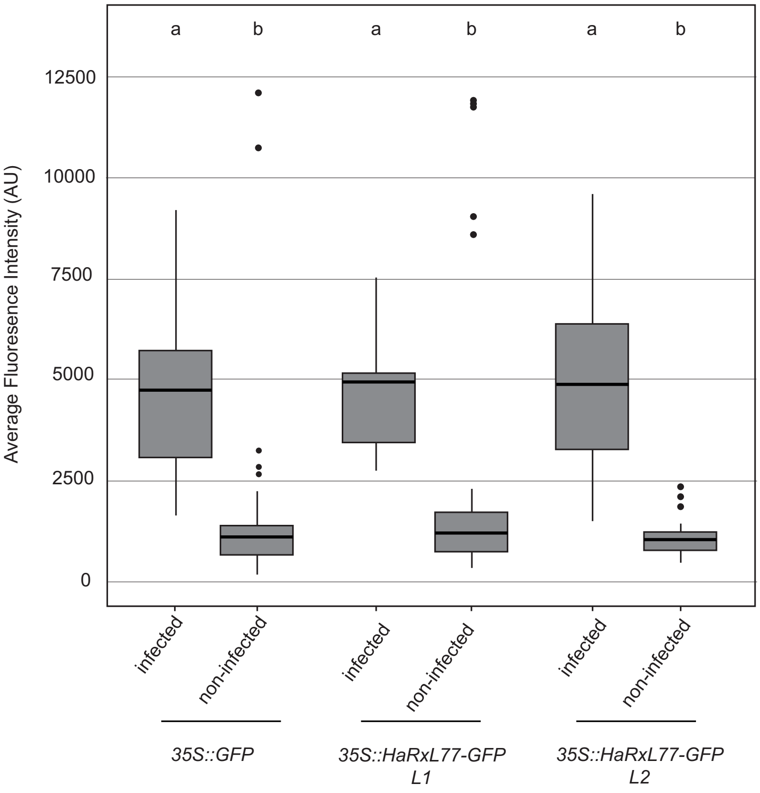


**Figure S6.** ***35S::HaRxL77-GFP*** **do not show excess callose deposition in the walls of infected cells.** 2-week-old seedlings were infected with *Hpa* waco9. Aniline blue stained leaves were imaged by confocal microscopy and the fluorescence intensity (indicative of callose abundance) was quantified by ImageJ. Data was analysed by ANOVA with Tukey post-hoc analysis for multiple comparisons, different letters indicate statistically significant differences ($p$ <0.005) . Box plots: The blue dot within the box marks the mean, the box signifies the upper and lower quartiles, and the minimum and maximum within 1.5 × interquartile range.
