## Supplemental Table 1 for "The *Hyaloperanospora arabidopsidis* effector HaRxL77 is hypermobile between cells and manipulates host defence"

Table S1. Screen of RxL effectors for mobility

| Highly expressed efffectors | Successfully synthesised | Localisation of effector | Mobility (H, hypermobile; M, mobile; R, reduced; I, immobile) |
|---|---|---|---|
| ATR13 | N | | |
| ATR39_HaRxL48 | Y | Cytoplasm&Nucleus | n.a. (low expression) |
| HaRxL100b | Y | Endoplasmic Reticulum | |
| HaRxL102 | Y | Cytoplasm&Nucleus | n.a. (low expression) |
| HaRxL102b | Y | no expression | |
| HaRxL108 | Y | Cytoplasm&Nucleus | I |
| HaRxL11 | N | | |
| HaRxL118 | Y | Endoplasmic Reticulum | |
| HaRxL12 | Y | Endoplasmic Reticulum | |
| HaRxL121 | Y | Cytoplasm&Nucleus | I |
| HaRxL13 | Y | Cytoplasm&Nucleus | H |
| HaRxL131b | Y | Cytoplasm&Nucleus | M |
| HaRxL131c | Y | no expression | |
| HaRxL135 | Y | Cytoplasm&Nucleus | M |
| HaRxL136c | Y | Cytoplasm&Nucleus | M |
| HaRxL14b | Y | Cytoplasm&Nucleus | H |
| HaRxL154 | Y | Chloroplast | |
| HaRxL25 | Y | Nucleus | |
| HaRxL25b | Y | Endoplasmic Reticulum | |
| HaRxL29 | Y | Endoplasmic Reticulum | |
| HaRxL34b | Y | Plasma membrane | |
| HaRxL35b | N | | |
| HaRxL37 | Y | Plasma membrane | |
| HaRxL37b | Y | Plasma membrane | |
| HaRxL37c | Y | Plasma membrane | |
| HaRxL37d | Y | Plasma membrane | |
| HaRxL44 | Y | Chloroplast | |
| HaRxL46 | Y | Cytoplasm&Nucleus | M |
| HaRxL57 | Y | Cytoplasm&Nucleus | H |
| HaRxL59 | Y | Plasma membrane | |
| HaRxL60 | Y | Chloroplast | |
| HaRxL61 | Y | Chloroplast | |
| HaRxL67 | N | | |
| HaRxL67b | N | | |
| HaRxL76 | Y | Endoplasmic Reticulum | |
| HaRxL77 | Y | Cytoplasm&Nucleus | H |
| HaRxL78 | Y | Cytoplasm&Nucleus | M |
| HaRxL8 | Y | Cytoplasm&Nucleus | H |

| | | | |
|---|---|---|---|
| HaRxL88 | Y | Chloroplast | |
| HaRxL92 | Y | no expression | |
| HaRxL94 | Y | Nucleus | |
| HaRxL94b | Y | Chloroplast | |
| HaRxL95 | Y | Nucleus | |
| HaRxL97 | Y | Cytoplasm&Nucleus | n.a. (stress symptoms) |
| HaRxL97b | Y | no expression | |
| HaRxL98 | Y | Endoplasmic Reticulum | |
| HaRxLCRN22 | Y | Cytoplasm&Nucleus | I |
| HaRxLCRN22b | Y | Cytoplasm&Nucleus | n.a. (low expression) |
| HaRxLL110 | Y | Cytoplasm&Nucleus | n.a. (low expression) |
| HaRxLL128b | N | | |
| HaRxLL137c | N | | |
| HaRxLL141 | Y | Endoplasmic Reticulum | |
| HaRxLL167 | Y | Cytoplasm&Nucleus | n.a. (stress symptoms) |
| HaRxLL169 | Y | Endoplasmic Reticulum | |
| HaRxLL169c | Y | Endoplasmic Reticulum | |
| HaRxLL21 | Y | Cytoplasm&Nucleus | H |
| HaRxLL29 | Y | Nucleus | |
| HaRxLL2b | Y | Nucleus | |
| HaRxLL38 | Y | Nucleus | |
| HaRxLL39b | Y | Nucleus | |
| HaRxLL427 | Y | no expression | |
| HaRxLL430 | Y | Endoplasmic Reticulum | |
| HaRxLL445 | Y | Cytoplasm&Nucleus | n.a. (stress symptoms) |
| HaRxLL449 | Y | Cytoplasm&Nucleus | H |
| HaRxLL455 | Y | Cytoplasm&Nucleus | n.a. (stress symptoms) |
| HaRxLL461b | Y | Endoplasmic Reticulum | |
| HaRxLL462 | Y | Cytoplasm&Nucleus | n.a. (stress symptoms) |
| HaRxLL464b | Y | Cytoplasm&Nucleus | n.a. (stress symptoms) |
| HaRxLL468 | Y | Cytoplasm&Nucleus | R |
| HaRxLL468b | Y | Cytoplasm&Nucleus | M |
| HaRxLL47 | Y | Nucleus | |
| HaRxLL470 | Y | Cytoplasm&Nucleus | H |
| HaRxLL5 | Y | Endoplasmic Reticulum | |
| HaRxLL50b | N | | |
| HaRxLL60 | Y | Cytoplasm&Nucleus | n.a. (stress symptoms) |
| HaRxLL62 | Y | Cytoplasm&Nucleus | M |
| HaRxLL62b | Y | Endoplasmic Reticulum | |
| HaRxLL65 | Y | Chloroplast | |
| HaRxLL67 | N | | |
| HaRxLL74b | Y | Cytoplasm&Nucleus | n.a. (low expression) |
| HaRxLL75c | Y | Cytoplasm&Nucleus | n.a. (low expression) |

| | | | |
|---|---|---|---|
| HaRxLL75d | Y | Cytoplasm&Nucleus | n.a. (low expression) |
| HaRxLL75e | Y | Endoplasmic Reticulum | |
| HaRxLL79b | Y | Cytoplasm&Nucleus | n.a. (low expression) |
| HaRxLL79c | N | | n.a. (low expression) |
| HaRxLL83 | Y | Cytoplasm&Nucleus | n.a. (low expression) |
| HaRxLL87 | Y | no expression | |

Notes: n.a.(stress symptoms) indicates no mobility screening was performed because the effector induced indicators of stress; n.a. (low expression) indicates mobility screening was impaired by low expression
